# Integrated analysis of crosslink-ligation data for the detection of RNA 2D/3D structures and interactions in vivo

**DOI:** 10.1101/2025.06.24.661343

**Authors:** Wilson H. Lee, Minjie Zhang, Zhipeng Lu

## Abstract

The latest advancements and implementation of crosslinking– and proximity-ligation based methods have transformed the study of RNA structure and interactions in living cells. Despite this progress, sequencing data must be meticulously processed to elucidate information on RNA. This presents distinct challenges, as noted in the Psoralen Analysis of RNA Interactions and Structures (PARIS) and Spatial 2′-Hydroxyl Acylation Reversible Crosslinking (SHARC) studies. In this chapter, we detail the process used to analyze RNA sequencing data derived from crosslink-ligation based experiments. We detail the strategies used to pre-process raw data, map to a reference genome, classify reads, and assemble duplex groups. Further, we show an example of analyzing structural data from the ribosomal RNA.

## 1. INTRODUCTION

As one of the principal informational biomolecules, RNA is a fundamental component to various macromolecular machinery that enable normal cellular process [1, 2]. Over the last few decades, the classical belief of RNA functioning as merely a mediator between DNA genetic information and functional protein products has been replaced by the understanding that RNA is a versatile biomolecule with diverse functions [3-6]. RNAs not only engage in various aspects of gene regulation, protein synthesis, and post-transcriptional modifications, but have also been found to play dynamic roles in cell signaling and other intracellular interactions [7-10]. For instance, the multitude of non-coding RNAs (ncRNAs) in cells, including small and large RNAs have been shown to interact with other ncRNAs, messenger RNAs (mRNAs), and proteins [11-15].

Although the complexity of RNA is increasingly appreciated, the structures and interactions of RNA on a cellular level remain poorly understood [8, 16]. The secondary and tertiary structures of many known RNAs are still in an early stage of characterization, although tools such as high-throughput sequencing and computational analysis have expedited the discovery of new RNAs [17, 18]. Manipulation and observation of RNA properties is critical to structure elucidation, although it has long been arduous and technically challenging. Physical methods such as X-ray crystallography, nuclear magnetic resonance (NMR) spectroscopy, and cryo-electron microscopy (EM) have made substantial progress in solving 3D structures of RNAs and their associated complexes [19-23]. In parallel, computational approaches, such as calculations for minimum free energy conformations or conservation of structural motifs in various nucleotide sequences have been used to assist in nucleotide base-pairing and RNA secondary or tertiary structure prediction [24-26]. While these approaches yield structures of few small RNAs, they remain time consuming and unsuitable for in vivo analyses due to various sample type limitations (e.g., size, flexibility, shape, etc.) [27, 28].

Efforts in the development of next generation sequencing-based RNA structure methods have profoundly increased the number of studies on in vivo RNAs [29]. These methods combine various enzymes and chemical agents, followed by sequencing in high throughput. Broadly, they can be grouped into either footprinting-or proximity ligation-based approaches.

Footprinting-based methods use probes to modify RNA bases, which are then captured by reverse transcription and read by sequencing and downstream analysis [30, 31]. Examples such as dimethyl sulfate (DMS) footprinting and 2′-hydroxyl acylation analyzed by primer extension (SHAPE) have been previously used to probe the flexibility and reactivity of individual nucleotides [32-34]. These chemical methods result in an indirect one-dimensional analysis, useful for guiding structure prediction and providing structural context to the nucleotides.

More recently, crosslinking-based methods (e.g., CLASH, COMRADES, LIGR-seq, PARIS, SPLASH, and SHARC) have been used to interrogate RNA features more directly, yielding information such as spatial proximity between fragments [8, 35-39]. These methods typically employ chemical reagents that react with interacting RNA base pairs. Upon fragmentation and purification/enrichment of crosslinked RNA fragments, the arm ends are enzymatically ligated and sequenced in high throughput to yield a chimeric molecule “read”, wherein segments come from distinct regions or entirely different molecules.

Although the advent of the crosslinking-based methods has deepened our understanding of RNA structures and interactions, there remain major challenges in analyzing the data as generated by these experimental techniques. Factors such as random fragmentation, complex crosslinking and proximity ligation orientation, and non-continuous reads convolute the data processing [40, 41]. To address these problems, it is important to standardize a pipeline for the systematic analysis of newly generated and previously published experimental data.

To date, there are myriad studies that have detailed the analysis of sequencing data derived from different DNA and RNA studies [29, 42, 43]. The applications of and commentary on the computational analysis of the crosslinking-ligation based data is typically limited to their respective studies [35, 36, 44]. As such, an integrated analytical approach that utilizes robust tools allows for a reproducible analysis across different sequencing data obtained in crosslink-ligation studies [40, 45]. In this chapter, we detail a stepwise pipeline for the analysis of crosslink-ligation derived experimental data using available software and scripts. We expect that sequencing data analyzed in a similar fashion will allow for insightful results, leading to a deeper understanding of RNA structures and interactions in living cells.

## 2. MATERIALS

This chapter outlines a systematic approach for handling data derived from RNA crosslink-ligation studies (*see* **Fig. 1A**), including sections for preprocessing, mapping to a reference, secondary structure analysis via CRSSANT (crosslinked RNA secondary structure analysis using network techniques), and further analysis. Steps from each section can be used independently and tailored to specific analytical purposes. Pre-processing, although described, is not strictly necessary for subsequent steps since data handling varies between library preparation approaches (i.e., ligation-based, tagmentation-based, amplicon based) and choice of sequencing platform (*see* **Fig. 1B**). All reads must be mapped to a reference genome prior to CRSSANT steps and subsequent analysis (*see* **Fig. 1C**). Familiarity with Linux/Unix environments, command line interface, NGS short read technologies, and basic python and shell scripting skills is recommended.

**Figure 1.**
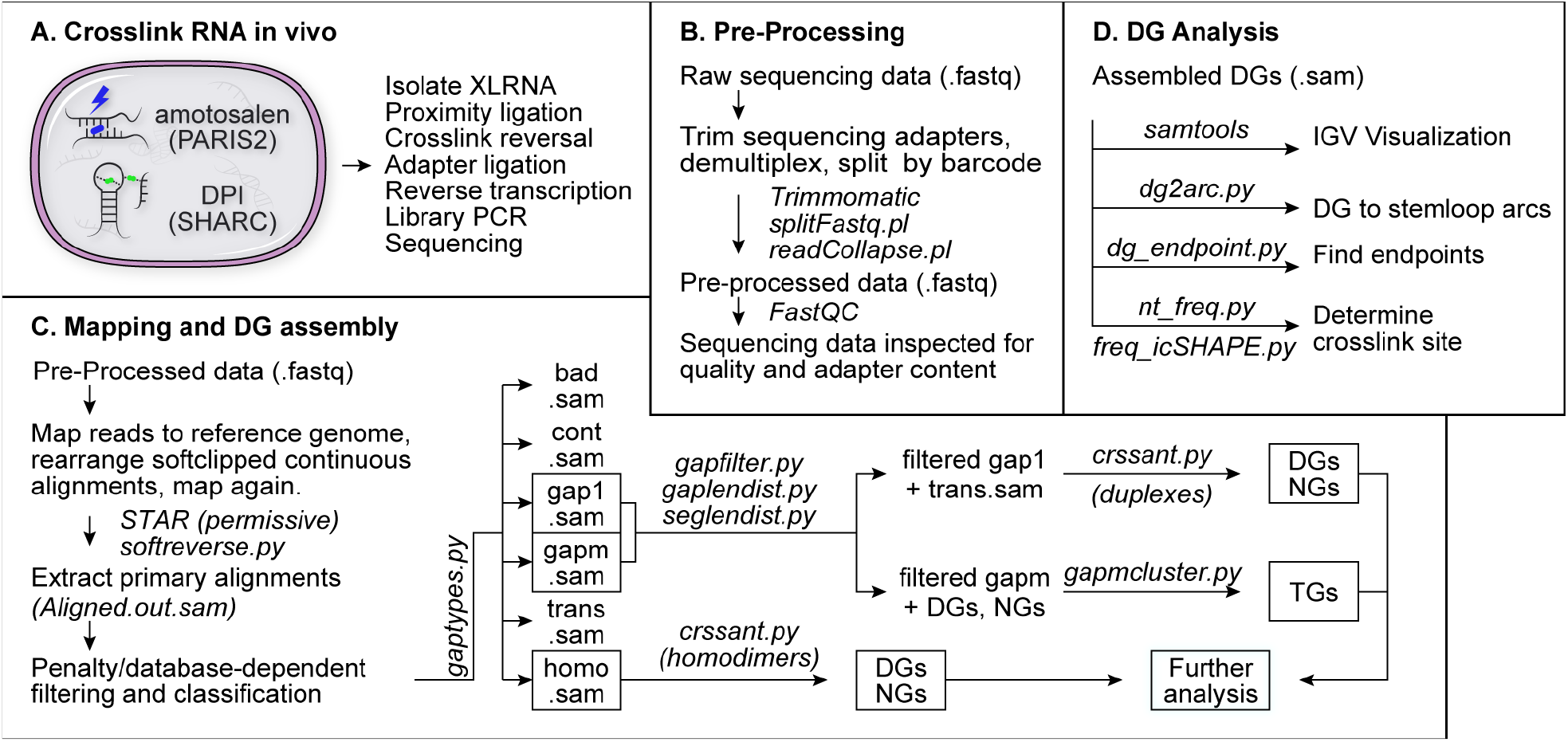
Stepwise diagram of the post-sequencing data handling process for crosslink-ligation based experiments. A) Overview of the crosslinking and library preparation steps leading up to sequencing. B) Pre-processing steps to clean up raw data, perform data quality control. C) Flowchart of the mapping and processes leading to CRSSANT analysis. D) Examples of further DG analysis.

### 2.1 Hardware Requirements

A high-performance compute (HPC) cluster with 64-bit Unix-based operating system, x86-64 compatible processer, and at least 32 GB of RAM is recommended for completing this protocol. Optionally, pre-processing steps can be performed locally on a Unix-based system prior to mapping and CRSSANT, both of which require an HPC cluster with >30 GB memory. Post-mapping and CRSSANT analysis steps can be performed locally on a system with at least 4 compute threads and 8 GB of RAM. Note that large datasets require more memory and upwards of one week for mapping and CRSSANT duplex group (DG) and non-overlapping group (NG) assembly. Disk space requirements are dependent on the size of raw sequencing reads (fastq) and subsequent aligned and processed read (sam/bam) files. For reference, operations described for pre-processing and mapping steps were performed on an HPC cluster (2.60 GHz 8-core Intel Xeon 2640v3, 32 GB, CentOS) and DG analysis steps on a standard desktop (3.3 GHz 4-core Intel Core i5, 8 GB, Ubuntu 20.04).

### 2.2 Software Requirements

As described, the protocol can be run entirely with the provided scripts and open-source tools. For detailed information regarding installation and troubleshooting, consult the manual, reference, or website linked with each tool (*see* **Note 1**). Use the software versions as detailed in this pipeline for error-free operation. While both legacy and updated versions of programs are available, various parameters and features may vary. Additionally, it is highly recommended that users have a GUI-based text editor to manipulate files and script parameters. Test datasets and example output files are supplied for crucial steps of the protocol (*see* **Note 2**). To allow for maximum parameter control and troubleshooting, each step of the pipeline should be run separately. Sample shell scripts and representative data for a typical pipeline are also provided as an example (*see* **Note 3**).

#### 2.2.1 Environment Setup

1. Python 3.x with packages biopython, matplotlib, NetworkX (v2.1+), numpy, pandas, pysam, scikit-learn, and SciPy. Note that Unix-based systems have a version of Python installed by default, but only versions 3.x+ will work with scripts in this protocol. See https://www.python.org/ for detailed installation instructions. We recommend using the Anaconda/Bioconda package managers to download and install dependencies.

#### 2.2.2 Pre-processing

1. Trimmomatic-0.36 [46]. http://www.usadellab.org/cms/uploads/supplementary/Trimmomatic/Trimmomatic-0.36.zip
2. icSHAPE pipeline [47]. https://github.com/qczhang/icSHAPE
3. FastQC [48]. https://www.bioinformatics.babraham.ac.uk/projects/fastqc/

#### 2.2.3 Mapping to Reference Genome

1. STAR-2.7.1a [49]. https://github.com/alexdobin/STAR https://github.com/alexdobin/STAR/archive/refs/tags/2.7.1a.tar.gz
2. samtools v1.1+ [50, 51]. https://www.htslib.org/download/ $ cd samtools-1.x $ make $ make install $ export PATH/bin:$PATH
3. Read classification and counting. https://github.com/whl-usc/lu_lab/archive/refs/

#### 2.2.4 CRSSANT and DG/NG Analysis

1. bedtools v2.22+ [52, 53]. https://github.com/arq5x/bedtools2
2. CRSSANT [40]. https://github.com/zhipenglu/CRSSANT/archive/refs/heads/master.zip
3. DG_analysis. https://github.com/whl-usc/rna2d3d
4. Integrative Genomics Viewer (IGV) [54, 55]. https://software.broadinstitute.org/software/igv/download

#### 2.2.5 Directory Setup and File Naming Conventions

1. Prepare directories for software, raw data, and working paths (*see* **Figure 2**). A directory containing relevant software (*see* **Figure 2A**) and references (*see* **Figure 2B**) will simplify access for future use (*see* **Note 4**). Pre-processed and downstream analysis files should be kept in directories separate from raw data (*see* **Figure 2C**).
2. Keep file naming conventions consistent between analyses of different datasets. This simplifies any changes to shell and python scripts when processing multiple datasets and prevents files from being erroneously overwritten. Table 1 shows an example of filenames with sample name/prefix “x”.

**Figure 2.**
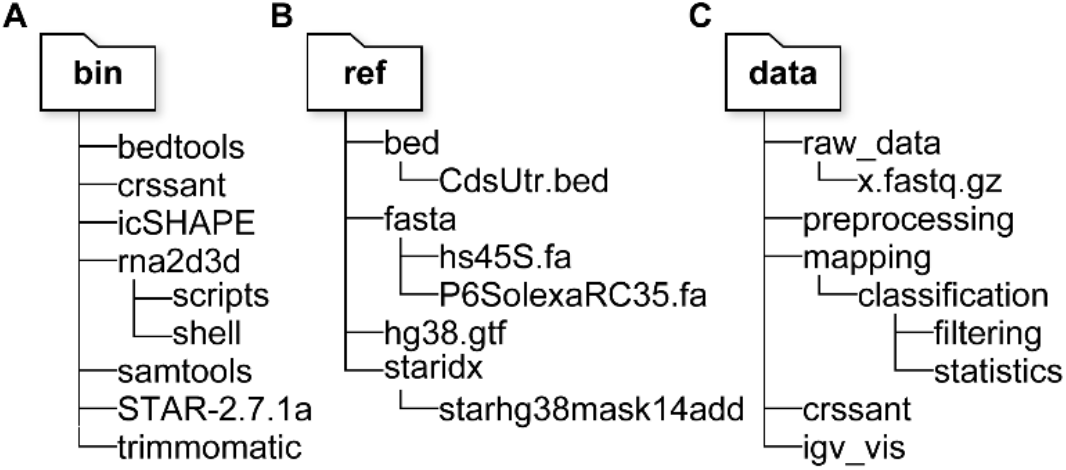
Sample directory trees for files relevant to the analytical pipeline. Note that only names of key files and sub-directories are included for clarity. Refer to the corresponding sections and Table 1 for further details. Consolidated directories contain the following: A) all relevant software and scripts B) reference sequences, genomes, and bed files C) data files, isolated into appropriate sub-directories.

**Table 1.** Example file names and associated descriptions. Note that the naming convention as described in the table assumes multiple barcodes (“bcnn.fastq”) are used and that demultiplexing is performed on the sample(s).

| File Names | Description |
| --- | --- |
| *Pre-processing (Trimming, demultiplexing, splitting by barcode)* |  |
| x.fastq.gz | Raw data (compressed) |
| x_trim3.fastq | 3'-end adapter sequence removed |
| bcnn.fastq | Split library based on barcode and name (e.g., CGTGAT:RTR701 has “bcnn” RTR701) |
| x_trim3_nodup.fastq | PCR duplicates removed |
| x_trim_nodup_bcnn.fastq | 5'-end adapter sequence removed |
| *STAR mapping to genome (permissive)* |  |
| x_Aligned.sortedByCoord.out.bam | All aligned reads mapped to STAR index, sorted by coordinate |
| x_Aligned.out.bam | All aligned reads mapped to STAR index, unsorted and in BAM format |
| xLog.final.out | Overall summary of mapping statistics |
| xLog.out | Main log file, detailed run information |
| xLog.progress.out | Job progress statistics (e.g., processed read number, mapped read statistics, etc.) |
| SJ.out.tab | High confidence splice junctions |
| Unmapped.out.mate1 | Output of unmapped or partially mapped reads |
| *Extracting primary alignments* |  |
| x_nonchimeric_temp.sam | Reads not aligned to multiple strands or chromosomes |
| x_chimeric_temp.sam | Reads aligned to multiple strands or chromosomes |
| x_nonchimeric_pri.bam | Non-chimeric primary reads converted to BAM format, discards supplementary and secondary reads |
| x_nonchimeric_pri.sam | Non-chimeric primary reads converted to SAM format |
| x_pri.sam | Combined primary non-chimeric and chimeric reads |
| *Classifying alignments* |  |
| x_cont.sam | Continuous alignments, no gaps |
| x_gap1.sam | Non-continuous alignments, 1 gap |
| x_gapm.sam | Non-continuous alignments, >1 gap |
| x_trans.sam | Non-continuous alignments, 2 arms on different strands or chromosomes |
| x_homo.sam | Non-continuous alignments, 2 arms overlap each other |
| x_bad.sam | Non-continuous alignments, complex combinations of indels and gaps |
| x_log.out | Log file for the run, details input and output alignment count |
| *Filtering spliced and short gaps* |  |
| x_prigap1_filtered.sam | x_prigap1 filtered for splice junctions, short deletions |
| x_prigapm_filtered.sam | x_prigapm filtered for splice junctions, short deletions |
| x_prigaps_filtered.sam | Combined prigap1 and prigapm files for segment and gap statistics |
| *Reads segment and gap statistics* |  |
| x_gaplen.pdf | PDF figure showing cumulative distribution histogram of the gap lengths |
| x_seglen.pdf | PDF figure showing cumulative distribution histogram of the segment lengths |
| x_pri_biotype.txt | Distribution of RNA into various types |
| x_pri_ReadCount.txt | Count of reads mapped to specific genes |
| x_pri_RPKM.txt | RPKM values for each gene |
| X_RPM.txt | RPM values for each gene |
| *CRSSANT* |  |
| x_pri_crssant.sam | Combined SAM file containing both gap1 and trans reads from gatypes.py |
| x_pri_crssant_TAG.sam | CRSSANT primary reads file tagged with unique identifier for separation after DG assembly |
| header | Sequencing file header from any SAM file associated with the same sample |
| x_pri_crssant.cliques.t_o0.2_dg.bedpe | bedpe output file listing all duplex groups formed |
| x_pri_crssant.cliques.t_o0.2.sam | SAM file containing alignments assigned to DGS, with DG and NG annotations |
| *Assembling DGs/NGs for rRNA* |  |
| x_prigap1_hs45S.tmp | Ribosomal RNA subunit hs45S extracted from all primary reads with 1 gap, with potentially SA tag present (column 21) |
| x_hs45S.sam | Ribosomal RNA subunit hs45S, SA removed, sequencing header added |
| x_hs45S_####.sam | Predefined number of randomly selected (e.g., “####”) hs45S reads. |
| *Data visualization with IGV* |  |
| x.bam | Compressed binary representation of SAM file containing all primary reads |
| x_sorted.bam | BAM file sorted by reference coordinates |
| x_sorted.bam.bai | BAM file index, stores BAM file position |

## 3. METHODS

### 3.1 Data preprocessing

In this section, post-sequencing data is pre-processed to remove sequencing adapters, short reads, PCR duplicates, and demultiplexed by barcode, if applicable (*see* **Note 5**). Upon completion of these steps, sample sequencing data is processed and suitable for mapping to a reference genome (*see* **Note 6**).

#### 3.1.1 Trimming, PCR duplicate removal, splitting by barcode

1 Run Trimmomatic in single ended mode to trim the 3’-end adapter sequences. Use functions “ILLUMINACLIP” and “SLIDINGWINDOW” with 16 threads and base quality encoding phred33. Specify the input, output, and adapter sequence files (*see* **Notes 6-7**).

~~~
$ java-jar trimmomatic-0.36.jar SE-threads 16-phred33 **x**.fastq.gz **x**.trim3.fastq ILLUMINACLIP:P6SolexaRC35.fa:3:20:10 SLIDINGWINDOW:4:20 MINLEN:18
~~~

2 Remove PCR duplicates by using built-in random hexamers using script readCollapse (*see* **Note 8**).

~~~
$ perl readCollapse **x**_trim3.fastq **x**_trim3_nodup.fastq
~~~

3 Where applicable, split the multiplexed library by barcodes using splitFastq.pl (*see* **Note 9**). Use parameter “-U” to indicate single ended reads for the input file, “-l” to specify a list of strings denoting different libraries, and “-b” to specify barcode position and length. Optionally include “-s” to generate simple library statistics.

~~~
$ perl splitFastq.pl-U **x**_trim3_nodup.fastq-l CGTGAT:R701-b 6:6-s
~~~

4 Run Trimmomatic in single ended mode to trim the 5’-end adapter sequences. Use function “HEADCROP”, with 16 threads, base quality encoding phred33. Specify input and output files.

~~~
$ java-jar trimmomatic-0.36.jar SE-threads 16-phred33 **bcnn**.fastq **x**_trim_nodup_bcnn.fastq HEADCROP:17 MINLEN:20
~~~

#### 3.1.2 FastQC

1 Visually inspect the quality of the pre-processed data using FastQC: “File” > “Open” (*see* **Figs. 3, 4**).

**Figure 3.**
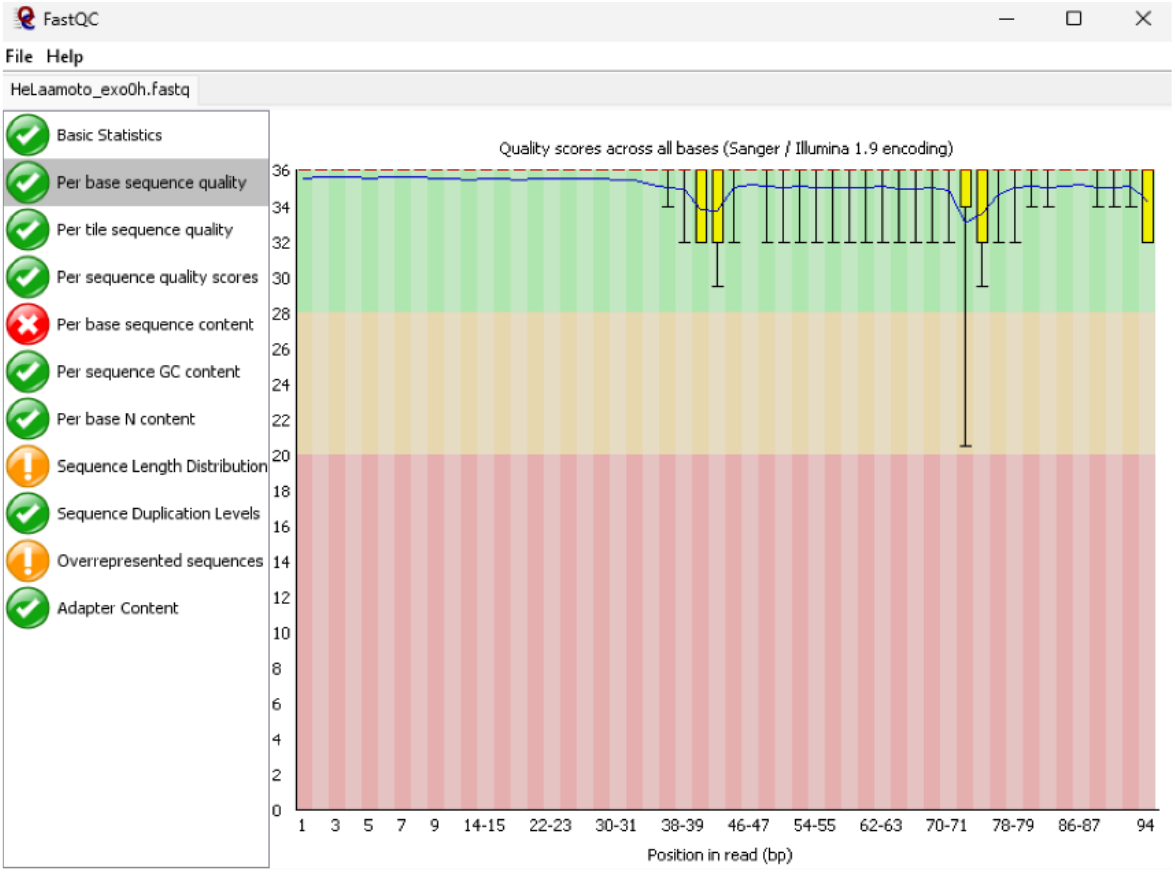
Screenshot of a fastq file loaded into FastQC. From the “per base sequence quality plot”, average quality score across all read positions should be high (>=30).

**Figure 4.**
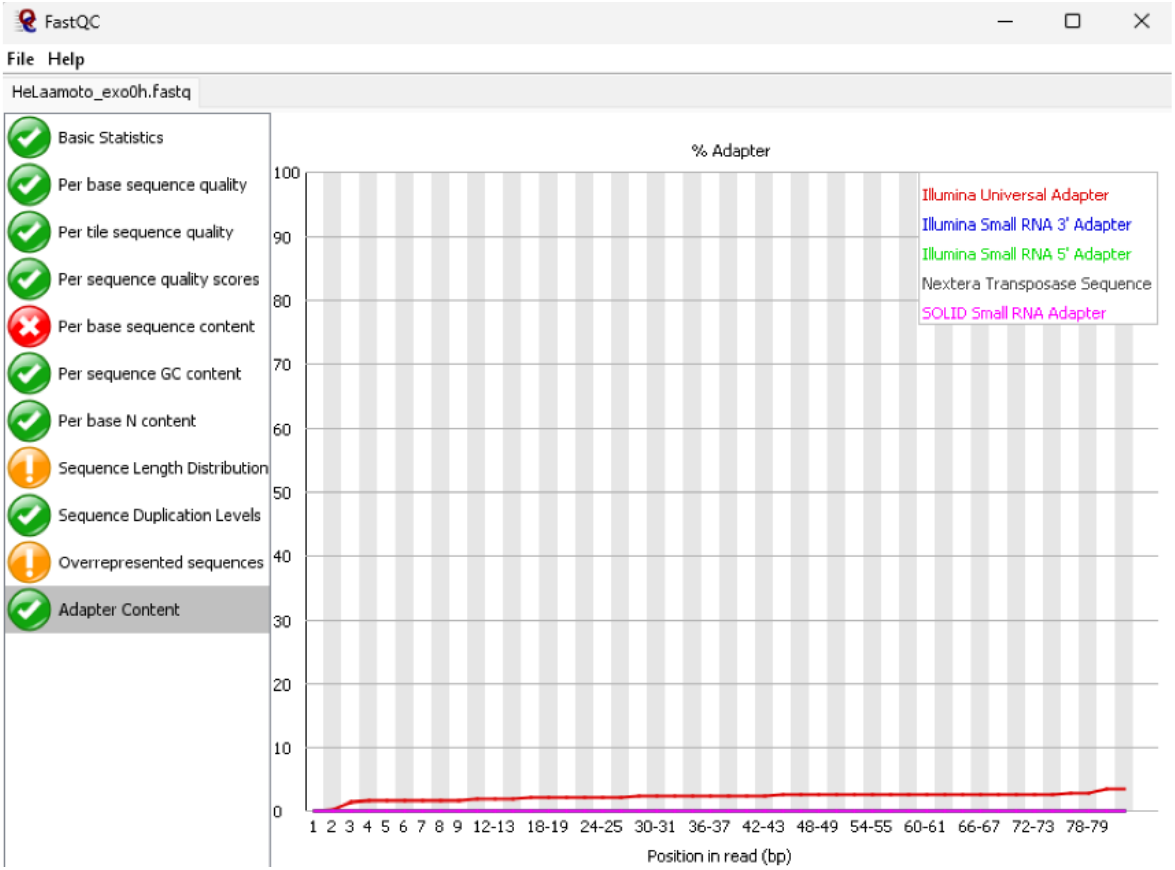
Screenshot of a fastq file loaded into FastQC. After sequencing adapter trimming steps, adapter content should be close to 0.

### 3.2 Mapping to reference genome, filtering reads, basic statistics

After data pre-processing, reads can be mapped to a reference genome of choice and classified into different alignment types, filtered for splice junction and short gapped reads, and plotted for gap and segment lengths.

#### 3.2.1 STAR mapping

1 Initialize STAR mapping to a reference genome with permissive parameters in a script (*see* **Notes 10-11**).

~~~
/STAR-2.7.1a/bin/Linux_x86_64/STAR \
--runThreadN 8 \
--runMode alignReads \
--genomeDir **StaridxPath** \
--readFilesIn **Fastq** \
--outFileNamePrefix **Outprefix**_1_ \
--genomeLoad NoSharedMemory \
--outReadsUnmapped Fastx \
--outFilterMultimapNmax 10 \
--outFilterScoreMinOverLread 0 \
--outSAMattributes All \
--outSAMtype BAM Unsorted SortedByCoordinate \
--alignIntronMin 1 \
--scoreGap 0 \
--scoreGapNoncan 0 \
--scoreGapGCAG 0 \
--scoreGapATAC 0 \
--scoreGenomicLengthLog2scale-1 \
--chimOutType WithinBAM HardClip \
--chimSegmentMin 5 \
--chimJunctionOverhangMin 5 \
--chimScoreJunctionNonGTAG 0 \
--chimScoreDropMax 80 \
--chimNonchimScoreDropMin 20
**Fastq** = **x**_trim_nodup_bcnn.fastq
**Outprefix** = **x**_1_
~~~

2 Extract primary alignments using samtools and awk.

~~~
$ samtools view-h **x**_Aligned.sortedByCoord.out.bam | awk ‘$1∼/^@/ || NF<21’ > x_nonchimeric_temp.sam
$ samtools view-h **x**_Aligned.sortedByCoord.out.bam | awk ‘$1!∼/^@/ && NF==21’> x_chimeric_temp.sam’
$ samtools view-bS-F 0×900-o **x**_nonchimeric_pri.bam **x**_nonchimeric_temp.sam
$ samtools view-h **x**_nonchimeric_pri.bam > **x**_nonchimeric_pri.sam
$ cat **x**_nonchimeric_pri.sam **x**_chimeric_temp.sam > **x**_1_pri.sam
$ rm-f **x**_nonchimeric_temp.sam **x**_chimeric_temp.sam **x**_nonchimeric_pri.bam **x**_nonchimeric_pri.sam
~~~

3 Classify primary reads based on alignment types using gaptypes.py (*see* **Notes 12-13**).

~~~
$ python3 gaptypes.py **x**_1_pri.sam **x**_1_pri-1 15 1
~~~

4 Rearrange any soft-clipped alignments using softreverse.py and map again using STAR with the same parameters as with above (*see* **Note 14**).

~~~
$ python softreverse.py **x**_1_pricont.sam softrev.fastq
/STAR-2.7.1a/bin/Linux_x86_64/STAR \
**Fastq** = softrev.fastq
**Outprefix** = **x**_2_
~~~

5 Extract primary alignments using samtools and awk again, using different file names.

~~~
$ samtools view-h **x**_2_Aligned.sortedByCoord.out.bam | awk ‘$1∼/^@/ || NF<21’ > x_2_nonchimeric_temp.sam
$ samtools view-h **x**_2_Aligned.sortedByCoord.out.bam | awk ‘$1!∼/^@/ && NF==21’> x_2_chimeric_temp.sam’
$ samtools view -bS -F 0×900 -o **x**_2_nonchimeric_pri.bam **x**_2_nonchimeric_temp.sam
$ samtools view -h **x**_2_nonchimeric_pri.bam > **x**_2_nonchimeric_pri.sam
$ cat **x**_2_nonchimeric_pri.sam **x**_2_chimeric_temp.sam > **x**_2_pri.sam
$ rm -f **x**_2_nonchimeric_temp.sam **x**_2_chimeric_temp.sam **x**_2_nonchimeric_pri.bam **x**_2_nonchimeric_pri.sam
~~~

6 Classify primary reads from the second round of mapping using gaptypes.py.

~~~
$ python3 gaptypes.py **x**_2_pri.sam **x**_2_pri -1 15 1
~~~

7 Combine the output file from both rounds of STAR mapping.

~~~
$ python3 merger_sams.py **x**_1_prigap1.sam **x**_2_prigap1.sam gap1
**x**_prigap1.tmp
$ python3 merger_sams.py **x**_1_prigapm.sam **x**_2_prigapm.sam gapm
**x**_prigapm.tmp
$ python3 merger_sams.py **x**_1_prihomo.sam **x**_2_prihomo.sam homo
**x**_prihomo.tmp
$ python3 merger_sams.py **x**_1_pritrans.sam **x**_2_pritrans.sam trans
**x**_pritrans.tmp
$ samtools view -H **x**_1_Aligned.sortedByCoord.out.bam > header
$ cat header **x**_prigap1.tmp > **x**_prigap1.sam
$ cat header **x**_prigapm.tmp > **x**_prigapm.sam
$ cat header **x**_prihomo.tmp > **x**_prihomo.sam
$ cat header **x**_pritrans.tmp > **x**_pritrans.sam
$ rm -f header *.tmp
~~~

#### 3.2.2 Filtering splices and short gaps

8. Remove all splice junction alignments and short deletions (1-2 nt gaps) from “gap1” and “gapm” read files using gapfilter.py.

~~~
$ python gapfilter.py Gtf **x**_prigap1.sam **x**_prigap1_filtered.sam 13 yes
$ python gapfilter.py Gtf **x**_prigapm.sam **x**_prigapm_filtered.sam 13 yes
~~~

### 3.2.3 Primary reads statistics (segments, gaps)

1 Plot the length of the gaps for filtered gap1 and gapm reads using gaplendist.py (*see* **Fig. 5a, Notes 15-16**).

~~~
$ cat **x**_prigap1_filtered.sam **x**_prigapm_filtered.sam >
**x**_prigaps_filtered.sam
$ python gaplendist.py **x**_prigaps_filtered.sam sam **x**.list all
$ python gaplendist.py **x**.list list **x**_gaplen.pdf all
$ rm -f **x**_prigaps_filtered.sam **x**.list
~~~

2 Plot the length of each segment and distribution for all types of gapped reads using seglendist.py (*see* **Fig. 5b, Notes 16-17**).

~~~
$ cat **x**_prigap1_filtered.sam **x**_prigapm_filtered.sam **x**_pritrans.sam > **x**_prifiltered.sam
$ python seglendist.py **x**_prifiltered.sam sam **x**_prifiltered.list
$ python seglendist.py **x**_prifiltered.list list **x**_seglen.pdf
$ rm -f **x**_prifiltered.sam **x**_prifiltered.list
~~~

3 Convert the primary reads file to BAM format. Determine the abundance of mapped reads to specific genes using script CountCdsUtr.py, which yields a summary of quantity and type of primary reads to the CDS, 5’-UTR, and 3’-UTR for genes in the hg38 genome (*see* **Notes 18-19**).

~~~
$ samtools view -bS -o **x**_prigap1_filtered.bam **x**_prigap1_filtered.sam
$ python CountCdsUtr.py **x**_prigap1_filtered.bam CdsUtr.bed none outname
Example: $ python CountCdsUtr.py test_prigap1_filtered.bam CdsUtr.bed none test
~~~

**Figure 5.**
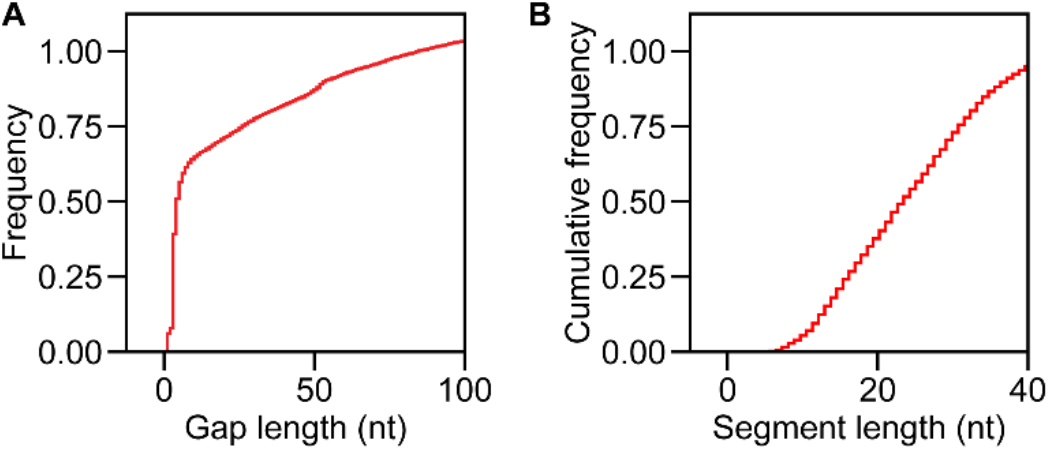
Example result a) gaplendist.py, distribution of “gap” lengths in STAR aligned reads b) seglendist.py, size distribution of “segment” length in STAR aligned reads.

**Figure 6.**
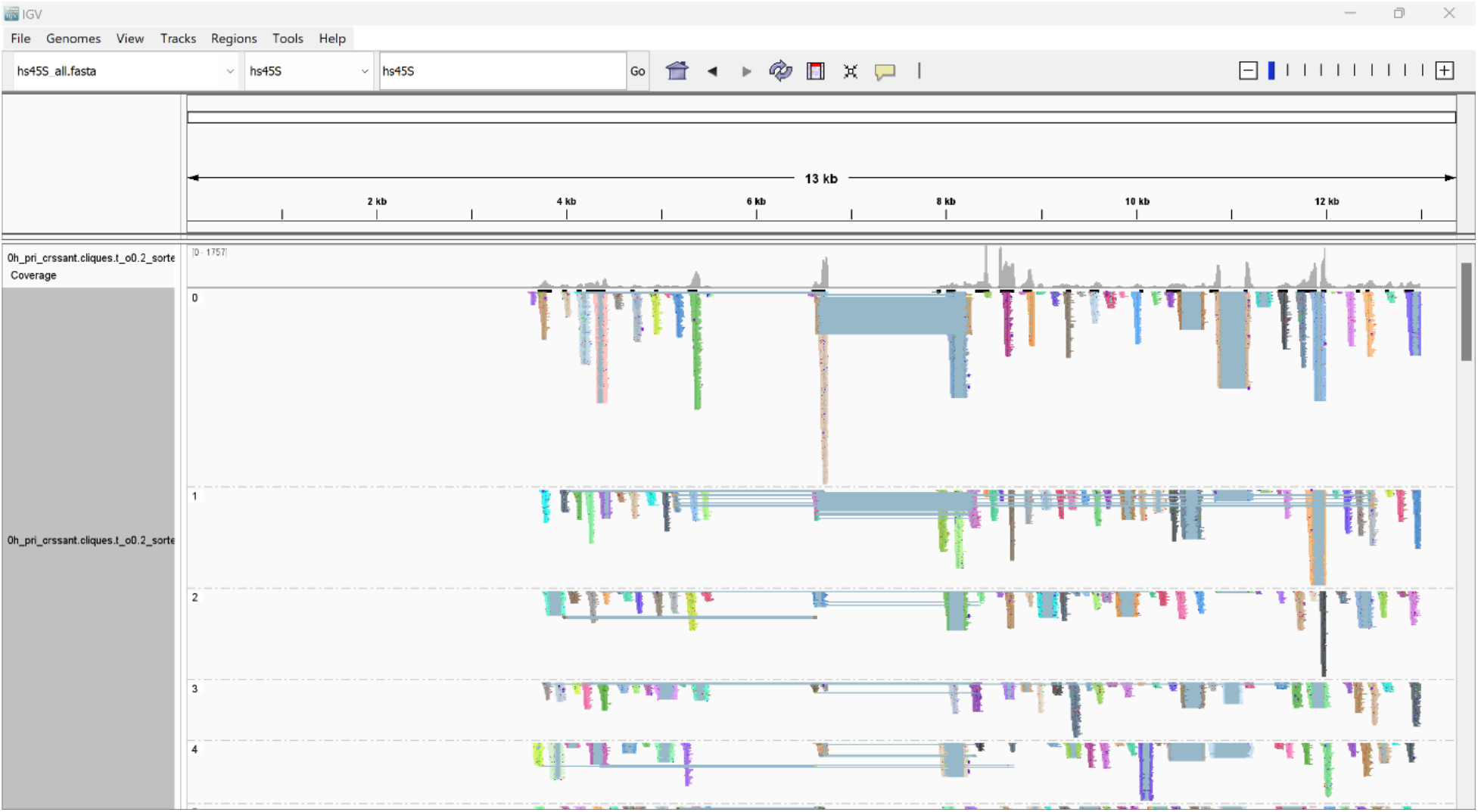
Screenshot of SAM file loaded into IGV with the hs45S rRNA subunit as reference genome. Visible here are the CRSSANT assembled reads from the hs45S subunit. The track is grouped by non-overlapping groups (NG), displayed in the “squished” setting.

**Figure 7.**
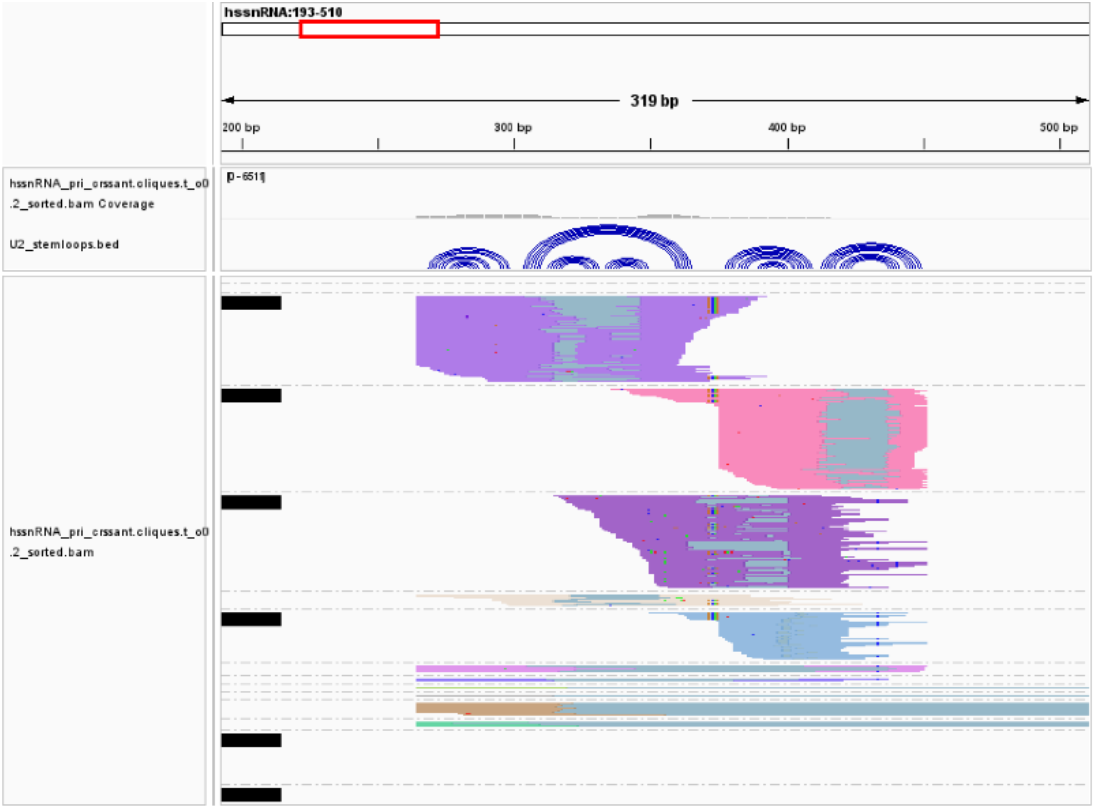
Example of using *dg2arc.py* to generate an arc file for U2 snRNA to show read arm interactions.

**Figure 8.**
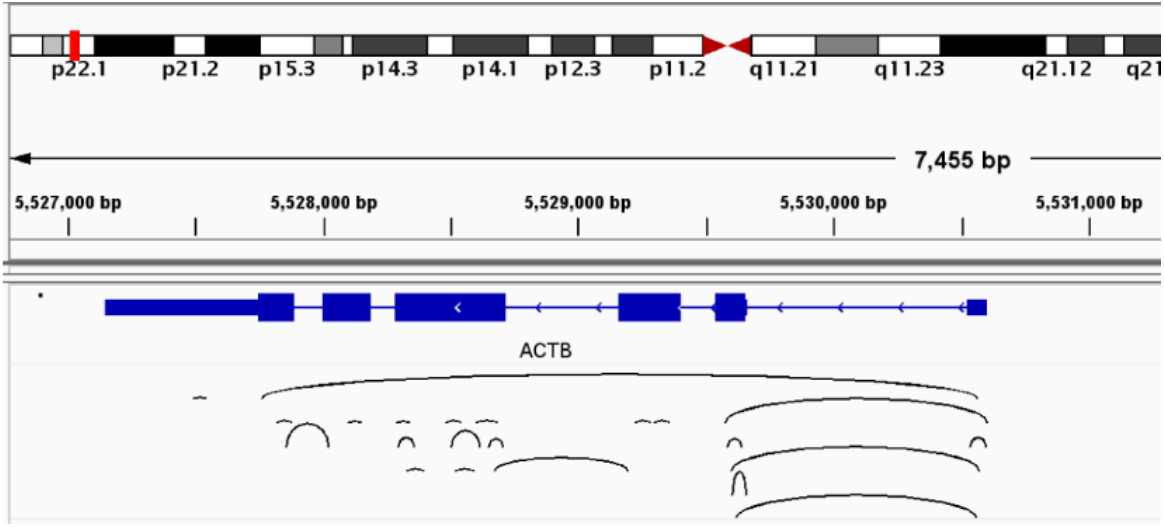
Example of using bedpetobed12.py to generate arc files for ACTB to show DG interactions.

**Figure 9.**
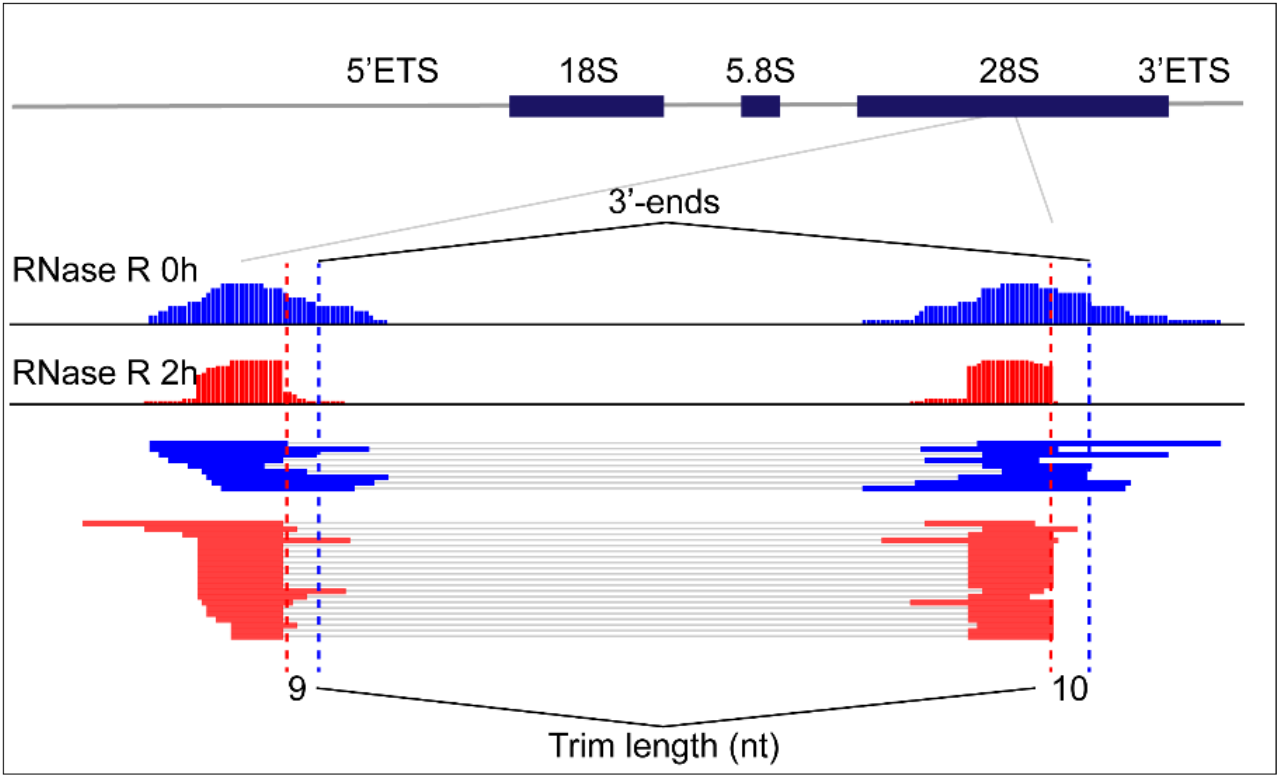
Example of the same CRSSANT generated DG from two sample types, with endpoints determined. Note the shortened read lengths in the RNase R treated “exo” (exonuclease) sample.

**Figure 10.**
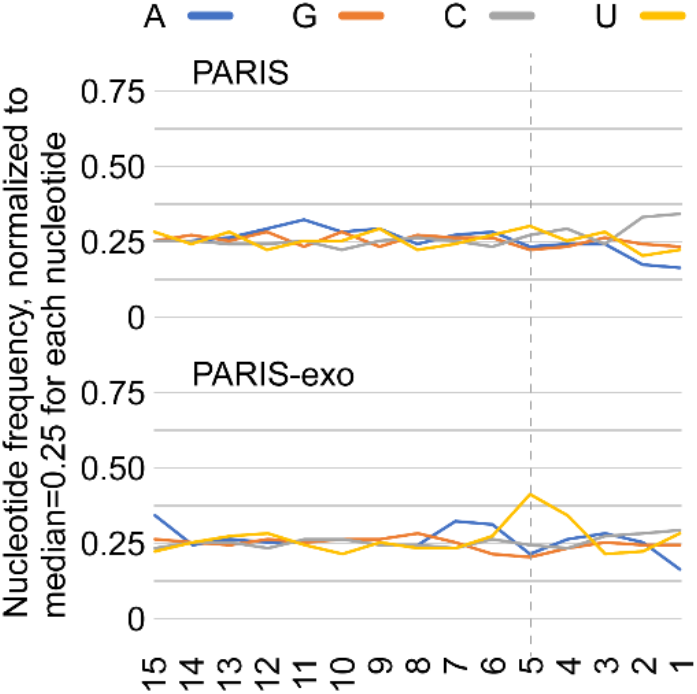
Example of two sets of data with nucleotide frequency determined for the 18S, 5.8S and 28S rRNA, 30,000 reads. Note the peak uridine frequency from the “exo” (exonuclease) treated sample.

**Figure 11.**
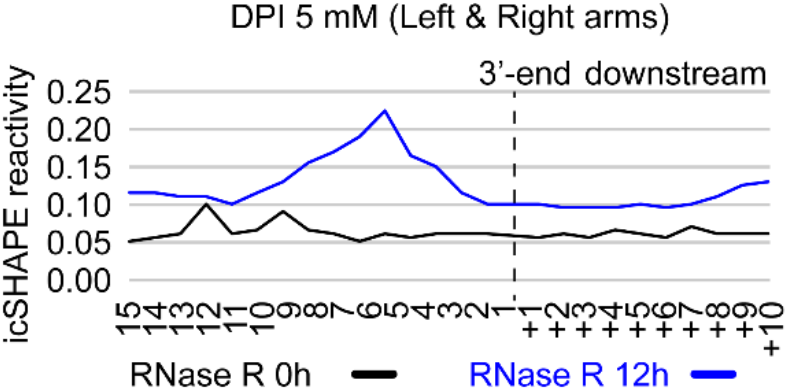
Example of two sets of data with SHAPE reactivity determined at positions -15 to 10 from the read arm ends. Note the sharp peak in reactivity 5 nucleotides from the arm end in RNase R treated sample.

## 3.3 CRSSANT – DG assembly and analysis

After STAR mapping, separation by gap type, and filtering, reads are output into distinct files as previously described. Depending on the end goal of the analysis, many options are available for further processing. For example, all types of gapped reads can be combined and used as input for CRSSANT analysis (i.e., duplex group (DG) and non-overlapping group (NG) assembly). Alternatively, reads mapped to a specific chromosome or ribosomal subunit can be extracted for quality control or comparison prior to DG or NG assembly. If performing CRSSANT immediately after mapping, modify relevant variables and paths and run DG assembly as part of the complete pipeline (*see* **Note 20**).

### 3.3.1 CRSSANT DG assembly

1 Use script merger.py to combine filtered gap1 and trans SAM files (*see* **Note 21**).

~~~
$ python merger_v2.py **x**_prigap1_filtered.sam **none x**_pri_crssant.sam
~~~

2 For different sample conditions, tag reads separately with a label according to experimental conditions, then combine the files with the sequencing header (*see* **Notes 22-23**).

~~~
$ awk ‘$0!∼/^@/ {printf $1”**-TAG1**\t”; for(i=2;i<=NF;++i) printf $i “\t”; printf”\n”}’ **x**_pri_crssant.sam > **x**_pri_crssant_**TAG1**_tmp.sam
$ awk ‘$0!∼/^@/ {printf $1”**-TAG2**\t”; for(i=2;i<=NF;++i) printf $i “\t”; printf”\n”}’ **x**_pri_crssant.sam > **x**_pri_crssant_**TAG2**_tmp.sam
$ samtools view -H **x**_pri_crssant.sam > header
$ cat header **x**_pri_crssant_**TAG1**_tmp.sam **x**_pri_crssant_**TAG2**_tmp.sam > **x**_pri_crssant.sam
$ rm -f header **x**_pri_crssant_**TAG1**_tmp.sam **x**_pri_crssant_**TAG2**_tmp.sam
~~~

3 Build a bed file for CRSSANT analysis (*see* **Note 24**). This can be done manually by following the tab delimited format described in the CRSSANT manual or by using genes2bed.py to process the “ReadCount.txt” output from CountCdsUtr.py.

~~~
$ python genes2bed.py **x**_ReadCount.txt CdsUtr.bed min_reads outname Example: $ python genes2bed.py test_ReadCount.txt CdsUtr.bed 10 min10
~~~

4 Use bedtools to generate a bedgraph for CRSSANT analysis.

~~~
$ bedtools genomecov -bg -split -strand + -ibam **x**_pri_crssant.bam –g **staridxPath**/chrNameLength.txt > **x**_plus.bedgraph
$ bedtools genomecov -bg -split -strand - -ibam **x**_pri_crssant.bam –g **staridxPath**/chrNameLength.txt > **x**_minus.bedgraph
~~~

5 Initialize CRSSANT for DG and NG assembly (*see* **Note 25**).

~~~
$ python crssant.py -cluster cliques -t_o 0.2 -out ./
**x**_pri_crssant.sam **Bed x**_plus.bedgraph,**x**_minus.bedgraph
Bed = bed file from step 3.
~~~

6 Cluster gapm alignments to tri-segment groups (TGs) using the output from the DG assembly.

~~~
$ python gapmcluster.py x.bedpe x_gapm.sam
~~~

7 Separate DGs by their tag after CRSSANT assembly is completed.

~~~
$ awk ‘$0∼/^@/ || $1∼/**-TAG**/’ **x**_pri_crssant.cliques.t_o0.2.sam > **x**_ crssant_TAG1.sam
~~~

### 3.3.2 Ribosomal RNA (rRNA) DG Assembly Example

Since rRNA reads are highly abundant, they are useful for comparative and quality control purposes. However, given the typically large number of rRNA reads, processing every read is unnecessary. Extracting reads mapped to the ribosome and using a randomized subset of them for further analysis is a convenient and recommended approach.

8. After mapping, extract only reads mapped to the ribosome and remove the SA tag (*see* **Notes 26-27**).

~~~
$ awk ‘$0∼/^@/ {print}; $0!∼/^@/ {if ($0∼”hs45S”) print}’
x_prigap1_filtered.sam > x_prigap1_hs45S.tmp
$ python3 merger_v2.py x_prigap1_hs45S.tmp none x_hs45S.sam
$ rm -f x_prigap1_hs45S.tmp
~~~

9 Extract a predefined number of randomly selected reads using extract_random.py (*see* **Notes 28-29**).

~~~
$ python extract_random.py **x**_hs45S.sam @ **30000 x**_hs45S_**30000**.sam
~~~

10. Repeat CRSSANT steps as previously described, this time using a bed file containing “genes” in the hs45S rRNA subunit (*see* **Note 30**).

~~~
$ samtools view -bS -o **x**_pri.bam **x**_hs45S_**30000**.sam
$ bedtools genomecov -bg -split -strand + -ibam **x**_hs45S_**30000**.bam -g
**staridxPath**/chrNameLength.txt > **x**_plus.bedgraph
$ bedtools genomecov -bg -split -strand - -ibam **x**_hs45S_**30000**.bam -g
**staridxPath**/chrNameLength.txt > **x**_minus.bedgraph
$ python crssant.py -cluster cliques -t_o 0.2 -out ./
**x**_hs45S_**30000**.sam **Bed x**_plus.bedgraph,**x**_minus.bedgraph
~~~

### 3.3.3 DG Analysis

Data produced after duplex group (DG) assembly via CRSSANT can be visualized with various tools.

1 Convert the CRSSANT output SAM file into BAM format, sort, and index. The samtools package is typically used in a three-step process as follows (*see* **Note 31-32**).

~~~
$ samtools view -bS -o **x**_pri.bam **x**_pri.sam
$ samtools sort –o **x**_pri_sorted.bam **x**_pri.bam
$ samtools index **x**_pri_sorted.bam
~~~

2 Open the BAM file in IGV, using the appropriate reference genome (*see* **Note 33**). Visualize the DGs, grouping by tag “DG” or “NG”.
3 Convert base pairs of DGs to arcs for IGV visualization (*see* **Note 34**).

~~~
$ python bedpetobed12.py x.cliques.t_o0.2_dg.bedpe
x.cliques.t_o0.2_dg.bed
$ sortBed -i x.cliques.t_o0.2_dg.bed > x.cliques.t_o0.2_dg_sorted.bed
Add header line track graphType=arc
~~~

4 Determine read coverage and DG arm endpoints (*see* **Note 35**).

~~~
$ python3 dg_endpoint.py input.sam
~~~

5 Count and plot the frequency of nucleotides at a given position from the 3’-end of reads (*see* **Notes 36-37**).

~~~
$ python nt_freq.py sam ref + - rRNA all_reads reads
**Example: $ python nt_freq.py x**_hs45S.sam hs45S.fasta 15 10 all n 30000
~~~

6 Count the frequency of SHAPE nucleotide reactivity at a given position from the 3’-end of reads. Useful for determining the crosslinking site of acylation-based experiments, efficacy of exonuclease trimming on the arm ends (*see* **Note 38**).

~~~
$ python SHAPE_freq.py inputsam shape_bedgraph DG_reads_cutoff DGcommon_ratio seqlen extendlen DG/reads outputprefix
Example: $ python SHAPE_freq.py 12h_pri_crssant.cliques.t_o0.2.sam HEK293_icSHAPE_hs45S.bedgraph 10 0.2 15 10 DG 12h_test
~~~

## 4. NOTES

1. Carefully read manuals of each tool and script. Detailed information on the execution of the scripts and programs, their options, and troubleshooting information is available online or within the script. For rna2d3d scripts, see usage instructions by running the python program with no arguments.
2. Relevant sample data to complete the protocol in its entirety can be downloaded from the original PARIS and CRSSANT references. For troubleshooting purposes, sample files are provided (<link>).
3. The pipeline allows for automation via shell scripting after completing any troubleshooting. Example shell scripts are provided (<link here>), all of which require modification of relevant variables and paths to run.
4. While not explicitly stated in each line of the protocol here, scripts, input, and output files should be given absolute directories or paths.
5. Each step can (and should be) initially be run independently. Depending on experimental design and method of cDNA library preparation prior to sequencing, pre-processing steps (and subsequently, the file names) may not necessarily be the same or omitted altogether.
6. The file size of raw sequencing data and number of barcodes passed through the script will affect the run time and output file sizes. For larger datasets, we highly recommended completing the steps on an HPC cluster.
7. For 3’-end trimming, parameters are as follow: seed_mismatches = 3, palindrome_clip_threshold = 20, simple_clip_threshold = 10, window_size = 4, window_minimum_quality = 20, minlen = 18. Ensure that the values are consistent with experimental design to not trim into sequencing reads.
8. After readCollapse, the printout to console will show total reads processed, unique number of reads after duplicate removal, and unique ratio (unique/total). This is a helpful set of numbers for basic quality control. Values can be kept in a spreadsheet for future reference and troubleshooting.
9. Multiple barcodes can be provided by using two colons “::” between “-l” barcode entries (e.g., CGTGAT:R701::ACATCG:R702). Whenever “-l” is used, barcode length “-b” must be specified. Files will be separated into fastq files based on the provided barcode name. Here, reads with barcode “CGTGAT” will be separated into output R701.fastq.
10. Settings for STAR mapping were previously optimized [41]. Briefly, it includes permissive parameters that enable sensitive and specific mapping of short reads with irregular gaps.
11. The parameters here are part of a shell script that initializes STAR mapping. The reference genome (StaridxPath), pre-processed data (Fastq), and output file prefix (Outprefix) must be specified.
12. Reads will be classified into 5 distinct primary types: cont (continuous, non-gapped), gap1 (1 gap, 2 segments), gapm (>1 gaps, >2 segments), trans (two segments, different strands/chromosomes, or homo (overlapping segments) reads. Maintaining a separate record of the file (i.e., sample origin, experimental conditions, etc.) will mitigate confusion over the details when comparing data, as files will have identical suffixes.
13. The original script “gaptypes.py” found in the CRSSANT repository is only compatible with Python2, so we recommend using gaptypes_v2.py (see rna2d3d repository) which is Python3 ready.
14. Details on the softreverse script were previously described [41]. Briefly, it accounts for discriminated backward chimeric alignments by rearranging softclipped continuous alignments from the first round of STAR mapping, which are then passed through a second round of STAR mapping.
15. The original script “gaplendist.py” found in the CRSSANT repository uses an older version of matplotlib, gaplendist_v2.py is updated to reflect these changes. However, we recommend using gaplendist_mod.py to simplify the plotting (see rna2d3d repository).
16. For files not filtered for splice junction alignments or short deletions, file input and output names would not include suffix “_filtered”. Subsequent file names would also omit the “filtered” suffix.
17. The original script “seglendist.py” found in the CRSSANT repository uses an older version of matplotlib, seglendist_v2.py is updated to reflect these changes. However, we recommend using seglendist_mod.py to simplify the plotting (see rna2d3d repository).
18. CdsUtr.bed is a bed file that specifies gene regions (CDS, intron, 5’-UTR, 3’-UTR) and type (protein coding, lncRNA, miRNA, rRNA, snRNA, etc.). Starting from a GTF file, only the longest isoforms of genes were used to manually prepare this bed file.
19. CountCdsUtr.py outputs various text files containing summarized information on the quantity and types of reads. For example, “_biotype.txt” calculates the percentage of mapped reads, while “_readCount.txt” details the frequency of reads mapped to a gene and its regions. Note that the CountCdsUtr script is only a rough estimate of read number, as it does not account for all isoforms. The example shown performs the read counting on gap1 reads.
20. The CRSSANT repository has been updated to include a python script that automates the pipeline. See the referenced page for details.
21. Replace argument pritrans.sam with any string for this step of analyses where only prigap1 or prigapm reads are used.
22. Tagging the reads is optional, particularly in cases where there is no separate dataset for comparison. However, it is highly recommended, as the reads will be given an identifier for future reference.
23. Substitute TAG with a short string that will distinguish it from other reads. This is especially important for experiments comparing similar data. For example, samples crosslinked at different concentrations might be given tags “0M” and “1M”.
24. The bed file here is tab separated with columns indicating chromosome, start and stop location, gene name, coverage for DG assembly, and strand. The script genes2bed.py uses the output “_readCount.txt” from CountCdsUtr to extract information for genes with a minimum read count greater than a given number (> “min_reads”) as cutoff.
25. One of the most common errors from CRSSANT is caused by the inclusion of the SA tag. To check for the SA tag, use grep or less and search column 21.
26. If necessary (i.e., comparing between a control and experimental sample), tag reads according to the conditions and re-add a sequencing header as previously described.
27. RNA reads were mapped to reference genome starhg38addmask14, which places the ribosomal RNA subunit hs45S into a separate “chromosome” named “hs45S”.
28. Here, number of reads = 30000. When comparing rRNA of two datasets, the number of reads should be limited by the dataset that has fewer reads. While this value can vary significantly between datasets, a good rule of thumb is to not exceed 5 M reads or use less than 0.01 M reads.
29. The SA tag was previously removed using merger.py. Here, since only gap1 reads are being processed for DG assembly, we must manually remove the tag, otherwise it will trigger an error when running CRSSANT.
30. This bed file contains the entire hs45S subunit, split into the “genes” 5.8S, 18S, 28S.
31. In this example, all primary reads are converted to BAM format. Oftentimes, converting, sorting, and indexing all mapped reads will take some time. Note that the CRSSANT output SAM file will have a suffix (i.e., “x_pri_crssant.cliques.t_o0.2.sam”).
32. The convert-sort-index process can be run line-by-line on a Unix/Linux system, using a shell script (see “samtools_igv.sh” in the rna2d3d repository), or simple python program (see “sam2bamsort.py” in the rna2d3d repository). The converted file can be visualized in software such as the Integrative Genomics Viewer (IGV).
33. BAM index files must be stored in the same directory as the BAM file being opened in IGV.
34. DGs must be on the same chromosome and strand to be converted into a bed12 file prior to visualization on IGV. Script dg2arc.py (see rna2d3d repository) is useful for manually curating arc-type interactions between read arms.
35. The output of “dg_endpoint.py” is a CSV file containing the DG number, coverage (how many reads in that DG) and the 50% average start/stop positions of each DG. By extracting the same information from two sets of relevant data, the DG read arm lengths can be compared. Note that this requires performing CRSSANT DG assembly for both sets of data, which would be combined after tagging with a unique identifier.
36. In the example here, the nucleotide frequency for x_hs45S.sam is determined, using the hs45S.fasta as reference. For 0 nucleotides downstream or “right” of the 3’-end, 15 nucleotides upstream or “left” of the 3’-end. All “genes” of the rRNA is selected, for “n” (not) all reads, but 30000 reads.
37. The output file is a CSV file containing the nucleotide position, and frequency of each nucleotide normalized to a median of 0.25. This information is especially relevant for determining the crosslinking site of psoralen- or formaldehyde-based experiments, as well as showing the relative efficiency of exonuclease trimming on read arm ends.
38. The output from “SHAPE_freq.py” displays the relative reactivity of nucleotide positions from upstream and downstream relative to the 3’-end of read arms. This is particularly useful in determining the expected crosslinking in acylation-based experiments and efficacy of exonuclease trimming on the arm ends.

## 5. ACKNOWLEDGEMENT

The authors thank both current and former lab members of the Lu lab for providing invaluable input and rigorous testing that have resulted in the culmination of this protocol.

